# Precision phase targeting of event-related oscillations using real-time closed-loop TMS-EEG

**DOI:** 10.64898/2026.03.26.713979

**Authors:** Malte R. Güth, Drew B. Headley, Travis E. Baker

**Affiliations:** Center for Molecular and Behavioral Neuroscience, Rutgers University, Newark, USA; Graduate Program in Neuroscience, Rutgers University, Newark, USA; Department of Biomedical Engineering, University of Minnesota, Minneapolis, MN, USA

**Keywords:** Real-time brain stimulation, Electroencephalography, Transcranial Magnetic Stimulation

## Abstract

**Objective:** Current closed-loop TMS-EEG systems rely on phase prediction algorithms that require highly periodic signals, limiting their ability to target brief, event-related activity. We developed a real-time closed-loop (RT-CL) TMS-EEG system that directly detects oscillatory phase without prediction, enabling phase-locked stimulation within microseconds.

**Methods:** We validated the system against a prediction-based approach using simulated sine waves and human EEG data (N=18), without active TMS delivery.

**Results:** Across frequency-modulated sweeps and spontaneous occipital alpha oscillations (eyes-open vs. closed), the RT-CL system achieved higher triggering probability (11-24%) and reduced the phase error variability (2-10°). Importantly, when targeting event-related theta oscillations during two spatial navigation tasks, RT-CL produced ∼20% higher triggering probabilities and ∼17° lower phase error variability than phase prediction.

**Conclusion:** These findings validate the RT-CL system for probing phase-dependent mechanisms during active cognition and pathological brain states.

**Significance:** By precisely targeting brief, variable neural signals, RT-CL could be used for the development of personalized TMS interventions for neuropsychiatric disorders during symptom provocation.

## I. Introduction

Transcranial magnetic stimulation (TMS) offers a unique combination of millimeter spatial resolution and millisecond temporal precision for non-invasively modulating neural activity in humans. This capability positions TMS as a tool for both probing causal relationships between neural oscillations and cognitive functions, and developing therapeutic interventions for neurological and psychiatric disorders [1], [2]. However, conventional open-loop TMS protocols deliver stimulation without regard to ongoing brain states, potentially missing optimal windows for neuromodulation. Closed-loop stimulation approaches address this limitation by triggering stimulation based on real-time EEG features, such as oscillatory phase, frequency, and power [3], [4], [5]. Accumulating evidence demonstrates that the responsiveness to TMS applied over the motor and visual cortex critically depends on when it is delivered relative to ongoing neural dynamics [6], [7], [8], thereby motivating the application of closed-loop approaches that synchronize stimulation with the phase of spontaneous EEG oscillations [9], [10], [11], [12].

To achieve phase-targeted stimulation with sufficient speed, current closed-loop approaches predict oscillatory phase from the preceding hundreds of milliseconds of EEG data, in some cases after calibrating parameters on prior baseline recordings [12], [13], [14], [15], [16], [17]. However, because prediction requires stable oscillatory properties, these methods are optimized for spontaneous oscillations with high periodicity but not for transient oscillations that emerge for only a few cycles following sensory and cognitive events (e.g., event-related potentials, phase-locked oscillatory activity). Moreover, EEG processing delays on PCs (e.g., data streaming, algorithm execution) and phase distortions (e.g., causal filtering) prevent real-time detection of oscillatory phase, forcing current closed-loop methods to rely on forward prediction of the phase [4], [18], [19]. This restricts both mechanistic investigation of phase-dependent neural processes and development of interventions targeting pathological brain states during active cognition.

To overcome these limitations, we developed a novel real-time closed-loop (RT-CL) EEG-TMS system using field programmable gate arrays (FPGAs), which enable microsecond-scale processing of multiple EEG channels in parallel and precise temporal control for phase-locked stimulation. This system was initially developed for animal studies in which optogenetic stimulation was triggered by amygdala gamma cycles during memory consolidation [20]. Importantly, this approach successfully modulated amygdala-dependent consolidation of spatial memory processes. Building on this, we adapted the system for non-invasive human application, enabling real-time detection and phase-locked TMS targeting of transient event-related oscillations (EROs) during cognitive tasks. To validate the RT-CL system, we compared its triggering probability and phase error with a phase prediction algorithm (PPA-CL) on simulated sine waves, spontaneous occipital alpha oscillations (eyes-open, eyes-closed), and EROs in the theta band during two spatial navigation reward tasks. To note, all validation was performed without TMS delivery to isolate system performance from stimulation artifacts. This validation study established RT-CL’s capacity to reliably detect and trigger TMS pulses to transient oscillatory signals during active cognition in real-time.

## II. Methods

### A. Experiment 1: Validation using Frequency-modulated chirps

To demonstrate the real-time FPGA system’s and prediction algorithm’s capabilities on a signal with a known ground truth, a function generator (AFG1062, Tektronix) was used to create a sinusoidal signal fed into the RT-CL and PPA-CL systems. Given the typical range of the EEG frequency spectrum and the frequencies of interest, a chirp signal descending in frequency from 50 Hz to 1 Hz lasting approximately 240 seconds was produced. The complete slow chirp was run once for peak, trough, ascending, and descending phase targeting. While running real-time triggering with the RT-CL system, these signals were saved to disk, so trigger timings could be analyzed and the chirp could be used for offline triggering with the PPA-CL system.

### B. Experiment 2: Validation using spontaneous alpha and event-related theta oscillations

#### Participants

For the human experiment, 25 undergraduate students (M_Age_ = 25 ± 2.76 years; 5 female) were recruited from Rutgers University Newark and the New Jersey Institute of Technology. Written informed consent was obtained from all participants. They were compensated with course credit as well as a monetary bonus equivalent to their memory performance in the T-maze and the Linear Track Memory (LTM) task. All subjects were screened with self-report questionnaires regarding demographic information, handedness [21], spatial navigation abilities [22], neurological and psychiatric history, their vision, and TMS permissibility (i.e., pregnancy, metal or medical implants, etc.). Only subjects who were right-handed, between 18 and 55 years of age, had normal-to-corrected vision, and had no history of neurological or psychiatric disorders or medication were allowed to participate. Three subjects were excluded due to incomplete data and three due to poor online EEG data quality, bringing the final sample to 19 subjects (M_age_ = 24 ± 2.51 years; 5 female). The study was approved by the Rutgers University Ethics Committee Board and was conducted in accordance with the ethical standards described in the 1964 Declaration of Helsinki.

#### Protocol

To induce spontaneous occipital alpha oscillations, participants were seated in a relaxed position in front of a black computer screen (Fig. 1A). Over the course of ten minutes, they underwent a series of alternating one-minute blocks of resting with their eyes open or closed. Each pair of conditions (eyes-open and eyes-closed) formed a two-minute cycle. A brief auditory cue was presented at the start of each 2.5-second transition interval to instruct participants to switch between the two states. This protocol was designed to elicit robust and alternating modulation of alpha-band activity over occipital regions [23]. The first block (one minute eyes-open, one minute eyes-closed) was used as a calibration period for the closed-loop targeting parameters (i.e., amplitude threshold). For the remaining four blocks, triggering alternated between ascending, peak, descending, and trough targeting. The order of phase targets was randomized and counterbalanced across subjects. Closed-loop triggering was focused on occipital alpha power (9-12 Hz) at the central occipital electrode Oz (Fig. 1A).

**Fig. 1.**
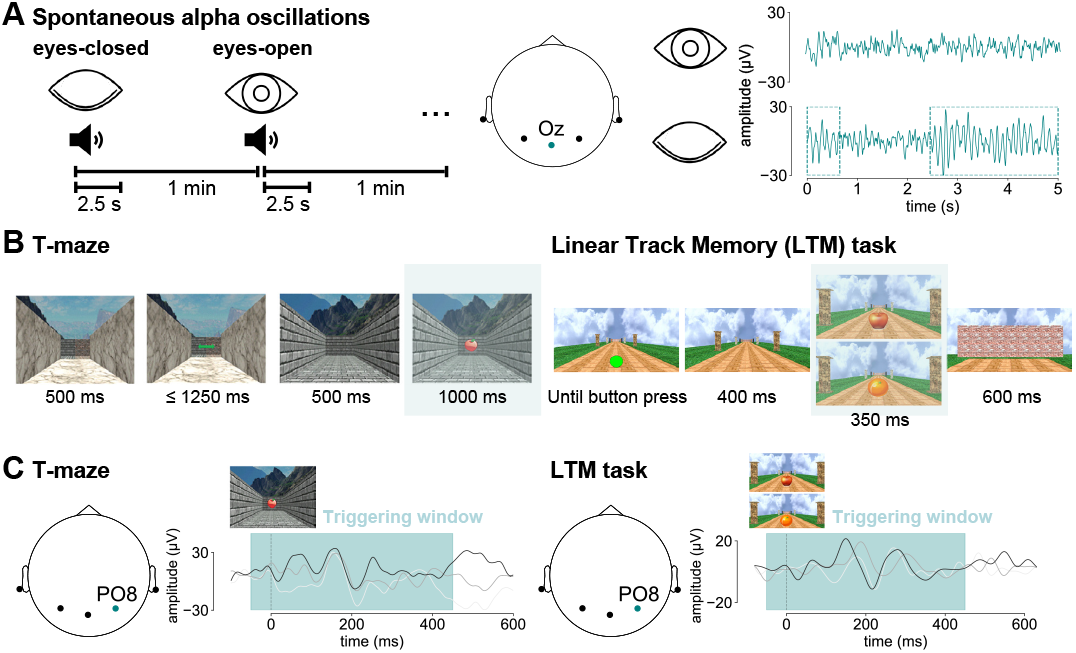
Tasks and triggering signals. **A**: Schematic of the paradigm for triggering on spontaneous alpha oscillations. Subjects alternated between closing and opening their eyes for one minute at a time. A tone indicated when to switch and triggering started 2.5 s after the onset of the tone. Triggering was performed on alpha oscillations recorded at Oz highlighted in teal on the 2D montage plot and the example traces separated by eyes-open (top) and eyes-closed (bottom). **B**: Overview of a single trial in the T-maze task (left) and LTM task (right). Teal rectangles highlight critical events for TMS triggering. On each trial subjects were placed in a T-shaped maze and chose to go left or right to search for rewards, which elicited RPT upon discovering feedback stimuli. On each trial of the LTM task subjects ran down a linear track towards a wall, passing five pairs of pillars where a reward (apple) or no reward cue (orange) appeared in the center of the track. **C**: Event-locked triggering time windows for the T-maze and LTM task. Triggering was performed on RPT recorded at channel PO8. The teal time window (-50 ms to 450 ms) on single-trial examples for trials from the respective tasks shows when triggering was enabled.

Subjects then performed two spatial navigation tasks used to elicit a burst of theta oscillations (4-10 Hz) over posterior electrodes, with larger amplitudes over right hemisphere channel PO8 [24], [25], [26]. This right posterior theta (RPT) response is characterized by spectral power peaking in the theta band around 200 ms following task-related feedback onset, and consistent with a partial phase reset of the oscillation [26]. First, participants performed 250 trials of a virtual T-maze task across four blocks (50 practice trials, 50 trials per block). In this task, participants navigated a virtual T-maze by choosing to go left or right in order to maximize monetary rewards (5 cents). Each trial began with a view down the stem of the maze toward a junction. After 1000 ms, a green double arrow appeared, prompting a choice: left (button 1, left index finger) or right (button 2, right index finger). Upon response, participants advanced to the chosen alley and, after 1000 ms, received visual feedback – an apple for reward or an orange for no reward – displayed for another 1000 ms. Feedback was randomly assigned with a 50% probability, unknown to participants (Fig. 1B, left). The practice trials were used as another calibration period to ensure that closed-loop triggering for RPT was set at an appropriate level for each subject. The four subsequent blocks were used to target the same four phase targets as before in a different counterbalanced order.

Following the T-maze task, subjects performed 120 trials of the LTM task (Fig. 1B, right, [27]) and 5 practice trials for calibration. Subjects were placed at the beginning of a straight track lined with uniquely colored and textured pillar pairs. On every trial, subjects started moving down the track after a green go sign appeared and they pressed a button to begin (400 ms ISI between pillars). At each pillar, subjects paused for 150 ms and a reward (apple) or no reward cues (orange) appeared for 350 ms in the middle of the track, each also eliciting RPT. At the end of the path, subjects stopped at a wall and were presented with a list of previously seen and distractor pillars. They were asked to indicate which cues they saw at a given pillar and could earn 5 cents per correctly identified reward cue location. For more details see Güth & Baker [27]. As before, closed-loop targeting was conducted in a counterbalanced order with one out of four phase targets being chosen for each block of 30 trials. In both the LTM and T-maze task triggering was enabled for from 50 ms before to 450 ms after the onset of feedback and reward cues respectively (Fig. 1C).

### C. Real-time closed-loop triggering

Real-time TMS-EEG synchronization is constrained by hardware- and software-related latencies [19], [28], necessitating systems to apply forward prediction and to estimate an expected delay from signal detection to TMS pulse delivery. These delays are highly dependent on the experimental setup and arise from multiple sources. Intrinsic delays due to the EEG data transfer to PCs (<1-2 ms [12], [29], [30]) or the output of a transistor-to-transistor logic pulse to trigger the TMS machine (1-2 ms [12], [15]) can be accounted for. More variable software-related delays such as data streaming with Lab Streaming Layer (∼6-12 ms [31]), TCP/IP (e.g., 50-100 ms [32]), or integrated acquisition and processing platforms like Open Ephys (total system delay ∼35 ms [33]) introduce latency jitters that differ across implementations (e.g., 10 ms jitter at 25 ms total delay [34]). Further, event-related, non-stationary signals like phase resetting can extend common prediction algorithms’ convergence period [17]. These variable delays can prevent accurate estimation of the total delay which has a substantial influence on the accuracy of the algorithm [18]. The RT-CL system addresses this limitation by processing analog signals directly on the FPGA, which enables parallel signal processing with microsecond-scale latency [20]. Unlike typical phase prediction implementations that process data through sequential stages, FPGAs perform all signal processing operations in dedicated hardware circuits. This eliminates the serial processing bottlenecks inherent to prediction-based approaches.

The closed-loop algorithm compiled on the FPGA was designed to track the amplitude and phase of EEG oscillations in real-time, acquire data, stream it to a PC for monitoring, and trigger TMS pulses. All programming was done in LabVIEW (National Instruments, Austin, TX, USA) and then compiled on a National Instruments USB-7845R reconfigurable data acquisition system based on a Xilinx Kintex-7 K70T FPGA. EEG data analysis, streaming, and acquisition was sampled at 1 kHz. While running, a fast direct memory access buffer between the PC and the FPGA was used to stream EEG signals for monitoring. Triggering was focused on channel Oz for alpha and PO8 for RPT, since in previous studies PO8 yielded the highest RPT power [24], [25], [26].

The triggering algorithm compiled on the FPGA consisted of four modules (Fig. 2, top): First, the analogue EEG signal was passed through a bank of Butterworth filters with 2^nd^ (for low pass) and 4^th^ (for high pass) orders (Fig. 2, module 1a). Each pair of filters acted as a band pass filter centered around a targeted frequency with a bandwidth of 2 Hz (alpha: 10 and 12 Hz; RPT: 4, 6, 8, and 10 Hz). Whichever filter band for amplitude detection had the highest power was selected at that moment. Applying multiple filters, and selecting between them in real-time, circumvented the problem of signal distortion as one moved away from the center frequency, as would be the case for a single bandpass filter encompassing these frequencies. In addition, the oscillation’s phase was analyzed using a second set of 1^st^ order Butterworth filters for the same bands, allowing for a more precise phase estimation (Fig. 2, module 1b). The lower order minimized phase distortion in the passband.

**Fig. 2.**
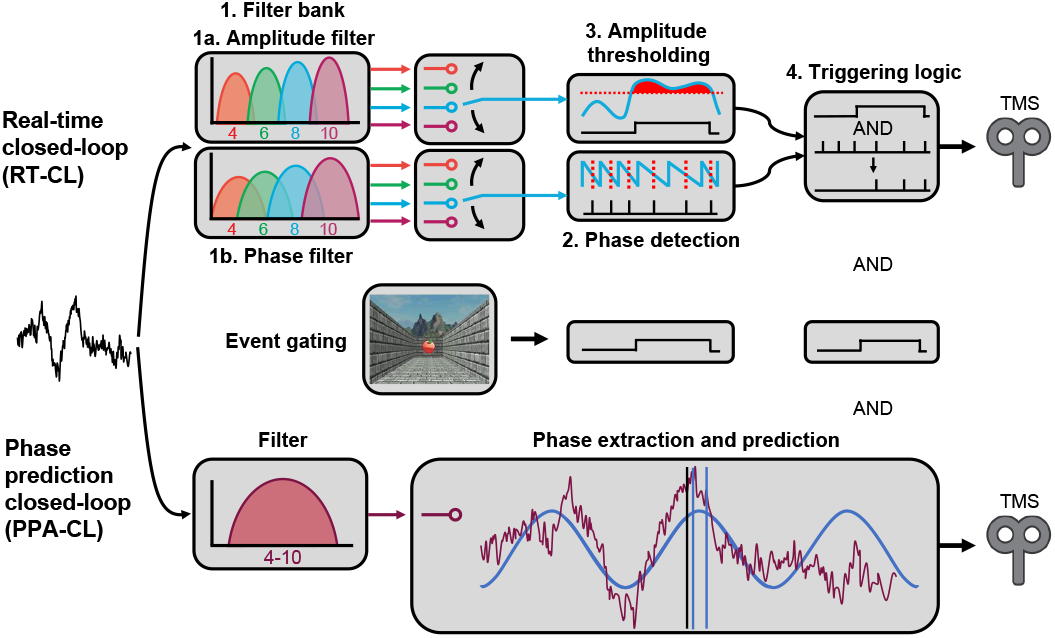
Closed-loop algorithms. The top row contains the RT-CL and the bottom row the PPA-CL. The middle row depicts the event gating logic, which was implemented in both algorithms. RT-CL phase analysis modules (from left to right): Bandpass filter bank centered around four theta frequencies for amplitude and phase estimation, phase detection, amplitude thresholding, and triggering logic. Triggering can be conditioned on the presence of event codes. The event gating mechanism used for the T-maze and LTM task added event code criteria and time constraint from a task like the feedback presentation. C: Single bandpass filter (9-12 Hz and 4-10 Hz) followed by fitting sinusoid waveform (blue) on Hilbert transform of EEG signal. Overlaid in red is the raw EEG signal. Triggering is enabled in crunch time window (blue vertical lines) on extrapolated sinusoid, reaching target criteria.

Second, peaks, troughs, descending, and ascending phases were identified based on sign changes of the signal waveform and its first derivative (Fig. 2, module 2). For each frequency band, the third module searched the first 5 ms after the occurrence of a peak or a trough for the maximum absolute value of the amplitude filter bank to assess the signal amplitude (Fig. 2, module 3). Third, the amplitude and phase of the strongest band were selected and sent to the last module for triggering. Picking the amplitude and phase of the strongest band ensured that the filter with the smallest phase error and the most pronounced signal was selected. The filter banks could be weighted to bias them towards detection in the theta or alpha bands. Fourth, the final module controlled whether a trigger was sent out after the targeted phase and frequency had been detected and the amplitude had exceeded the set threshold (Fig. 2, module 4). An event gating mechanism based on digital trigger inputs from the computer displaying the T-maze and LTM task ensured that a TMS pulse could only be triggered when a relevant task event was detected (i.e., feedback in the T-maze and reward cue in LTM task).

### D. EEG acquisition

The RT-CL system consists of FPGAs performing the signal processing and triggering, which require analogue EEG input signals. Contemporary EEG systems typically digitize signals and use USB protocols to transfer the data to a PC. Thus, an EEG8 amplifier (Contact Precision Instruments, Glastonbury, UK) was used to preserve the signal in its analogue form and to forward the data to the FPGA. EEG data were recorded from a custom montage of five channels including Afz as ground. Oz (for alpha) and PO8 (for RPT) served as trigger channels and were referenced online to the average of M1 and M2. PO7 was included as a backup channel. Data were acquired using Ag/AgCl ring electrodes mounted in a nylon cap with abrasive conductive gel (Falk Minow Services, Herrsching), sampled at 1 kHz, and sent directly to the FPGA. To reduce electrical noise, the amplifier was housed in a shielded container with the FPGA hardware. An online bandpass filter (0.1-100 Hz) and 60 Hz notch filter (50 dB/octave) were applied.

### E. Prediction-based triggering

To compare our approach to a conventional method for EEG phase-locked triggering, we used a phase prediction algorithm implemented in the EventIDE software environment (v. Feb 21, 2021, Okazo Lab, Achterom, Netherlands) to detect different phases of alpha and theta activity offline after data were recorded. A Fast Fourier Transform was continuously applied to the signal to estimate frequency spectra. This algorithm performed phase-locked triggering by fitting a sine wave onto the Hilbert transform of a buffered bandpass filtered EEG signal (alpha: high-pass: 9 Hz, low-pass: 12 Hz; RPT: high-pass: 4 Hz, low-pass: 10 Hz) and by extrapolating the signal into the future (Fig. 2, bottom). The forecast window was set to 200 ms and triggers for the same phase targets as for the RT-CL were sent within a crunch time window of 26 ms after the current sample.

The same calibration periods were used to set the amplitude threshold, the maximum model prediction error, and other triggering parameters. When adjusting parameters in either closed-loop system, this adjustment can be made in favor of sensitivity (i.e., more but less accurate triggering) or selectivity (i.e., less but more accurate triggering). To ensure comparability between the two closed-loop runs, we attempted to match the level of sensitivity by adjusting parameters for the offline PPA-CL runs until the number of pulses was similar to the number of pulses in the RT-CL run. These included the power threshold and the minimum model fit between the sine wave and the analytic signal.

### F. Analysis of closed-loop triggering performance

For preprocessing of the human EEG data, the respective triggering time windows were extracted. The signal was bandpass filtered with a causal second-order Butterworth filter in the targeted frequency range (alpha: 9-12 Hz, RPT: 4-10 Hz) for the instantaneous frequency estimation and with a first-order Butterworth filter in a wider range (alpha: 5-16 Hz, RPT: 1-14 Hz) for the instantaneous phase estimation. All stimulated phase angles and all true alpha and theta cycles exceeding the amplitude threshold used during the respective sessions were then extracted. Two parameters were analyzed to evaluate triggering performance for each system and subject: the triggering probability (i.e., sensitivity; Fig. 3A) and the standard deviation of the phase errors (SDPE, i.e., precision; Fig. 3B). Triggering probability was calculated by dividing the number of correctly detected phase targets of a given frequency by all true cycles of that frequency and phase target, yielding the probability of detecting and triggering on a single cycle. Phase error statistics were calculated as the circular mean and standard deviation of the difference between the targeted phase angle and the stimulated phase angles across trials.

**Fig. 3.**
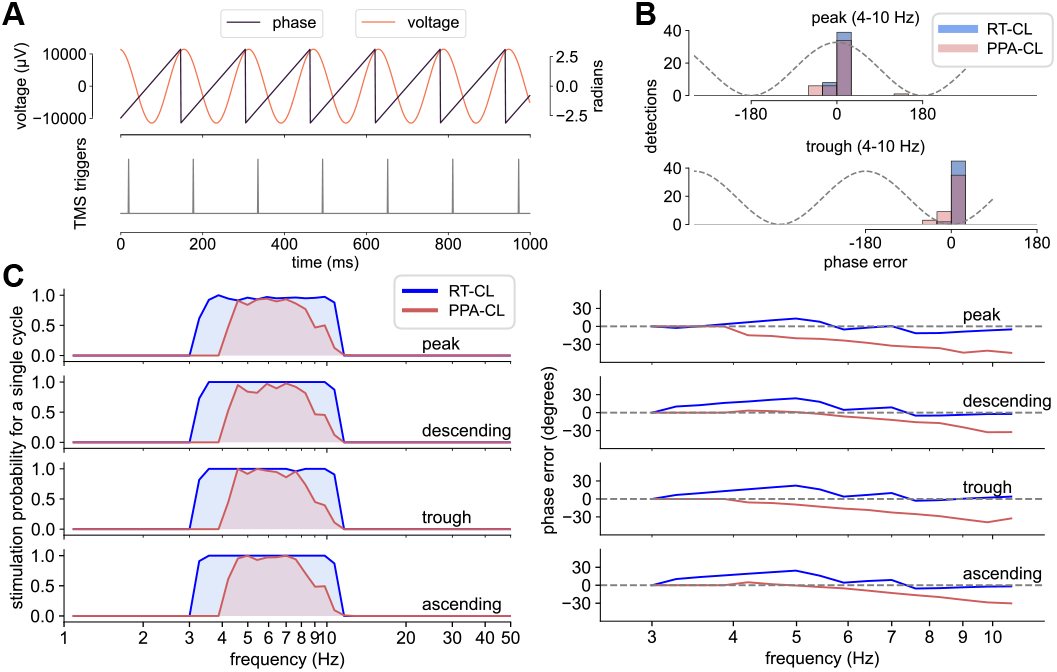
Triggering performance on simulated slow chirps. **A**: Schematic of theta detection and triggering, showing six simulated theta cycles (orange), their instantaneous phase angle (black), and the timing of the TMS triggers targeting troughs. Triggering probability was calculated as the ratio of real phase targets followed by a TMS trigger to the total number of real targeted phases. **B**: Phase timings of triggers sent with the RT-CL (blue) and PPA-CL (red) system targeting troughs and peaks overlaid on a theta sine wave. Phase angles are expressed as deviations from the targeted phase in degrees. The phase error was calculated as the standard deviation of phase timings across triggers. **C**: Right plot depicts triggering probability on a single cycle for RT-CL (blue) and PPA-CL (red) systems on the slow chirp (1-50 Hz) for each frequency target. Left plot shows the phase error across log-scaled frequency bins in degrees for RT-CL (blue) and PPA-CL (red) systems and for each phase target. Targets are plotted in the same row as on the left.

For the simulated chirp (Experiment 1), precision was assessed as the standard deviation across frequencies of the theta band, since each specific frequency between 1-50 Hz only occurred once. For the human EEG data (Experiment 2), the circular mean and standard deviation of the phase error while targeting theta and alpha were computed across trials for each subject. Performance differences between systems in triggering probability were assessed using two-tailed Wilcoxon signed-rank tests for Experiment 1 (treating individual theta frequencies as observations) and two-tailed paired t-tests for Experiment 2 (using individual subject averages as observations).

## III. Results

### A. Experiment 1: Frequency-modulated chirp

For RT-CL, triggering probability remained consistently high across the targeted theta band, with ∼100% triggering probability from 3.5-4 Hz until 10 Hz for every phase target (Fig. 3C, left). On average, a single theta cycle was triggered with a probability of 90.6% for peak targets, 91.7% for troughs, 91.3% for the descending phase, and 92.7% for the ascending phase. The slight deviations from 100% triggering probability occurred only outside the targeted range at the lower bound of the 4 Hz filter. When disregarding bins below 4 Hz, average triggering probabilities reached 98.6% (peak: 95.1%, trough: 99.6%, descending: 100%, ascending: 100%). In contrast, the PPA-CL system reached 100% triggering probability only around 5 Hz (92% at 4 Hz), with consistent performance in the 5-7 Hz range (91-94% triggering probability). However, at the lower and upper edges of the desired range (4 Hz and 10 Hz), the probability of stimulating a given cycle dropped substantially, falling below 10% at the upper edge of 10 Hz. Averaged across the theta frequency range, the PPA-CL system achieved triggering probabilities of 73.2% for peaks, 75.6% for troughs, 74.5% for the descending phase, and 77.2% for the ascending phase. A Wilcoxon signed-rank test comparing the average triggering sensitivity across phase targets between RT-CL and PPA-CL revealed significantly higher sensitivity for RT-CL (*W* = 0, *z* = -2.93, *p* = .003, *r*_*rb*_ = 1).

For phase precision, the SDPE of the RT-CL and PPA-CL systems was calculated (Fig. 3C, right). The RT-CL system demonstrated low variability in phase error, with SDPEs of 7° for peaks, 7.5° for troughs, 9.7° for the descending phase, and 9.7° for the ascending phase. This variability remained relatively unaffected by frequency, staying consistent across the theta range. Triggering with PPA-CL yielded slightly higher variability: 9.4° for peaks, 10.1° for troughs, 11.6° for the descending phase, and 10.7° for the ascending phase. This larger phase error stemmed from a frequency-dependent increase in stimulated phase, resulting in higher phase variability for the PPA-CL system with a variability ratio of 1.59 (PPA-CL divided by RT-CL). Comparing average SDPEs across phase targets, the RT-CL system achieved 19% higher precision than the PPA-CL system. Notably, both systems showed relatively consistent performance across phase targets (triggering probability: RT-CL 90.6-92.7%, PPA-CL 73.2-77.2%; SDPE: RT-CL 7-9.7°, PPA-CL 9.4-11.6°). Therefore, statistics for the Experiment 2 are presented for the average across phase targets. A complete table of performance measures by task, system, and phase target can be found in Supplementary Table 1.

### B. Experiment 2: Human EEG data

#### Alpha oscillations

Fig. 4A depicts an example trial targeting peaks of occipital alpha oscillations using RT-CL. Triggering occurred consistently within a few milliseconds of the occurrence of the phase target, demonstrating the system’s ability to detect and respond to spontaneous oscillatory activity in real-time. The two systems also differed in their timing relative to the targeted phase (Fig. 4B). The PPA-CL system delivered triggers before the targeted phase occurred, with an average phase error of -13° (negative values indicating early triggering), while the RT-CL system delivered triggers with an average phase error of 15.6°, occurring after the targeted phase as designed, *t*(18) = 4.54, *p* = 0.0002, *d* = 1.07. This timing difference is evident in the example trace (Fig. 4A) and in the circular histograms of phase errors across all subjects and phase targets (Fig. 4B). Visual inspection of these histograms confirmed narrower distributions for the RT-CL system across all phase targets, indicating more consistent triggering performance. For phase precision, measured as the SDPE, the RT-CL system demonstrated significantly lower variability than the PPA-CL system, *t*(18) = 8.27, *p* = 1.52 × 10^-7^, *d* = 1.95 (Fig. 4C, right). The RT-CL system showed an average SDPE of 29.2° (SD = 4.8°), compared to 39.6° (SD = 3°) for the PPA-CL system, representing approximately 10° higher precision. In terms on sensitivity, triggering probability was significantly higher for RT-CL than for PPA-CL, *t*(18) = 3.82, *p* = .001, *d* = 0.9 (Fig. 4C, left). Specifically, the RT-CL system detected approximately 70% of alpha cycles in the EEG signal (SD = 10.5%), compared to 59.1% (SD = 5.6%) detected by the PPA-CL system.

**Fig. 4.**
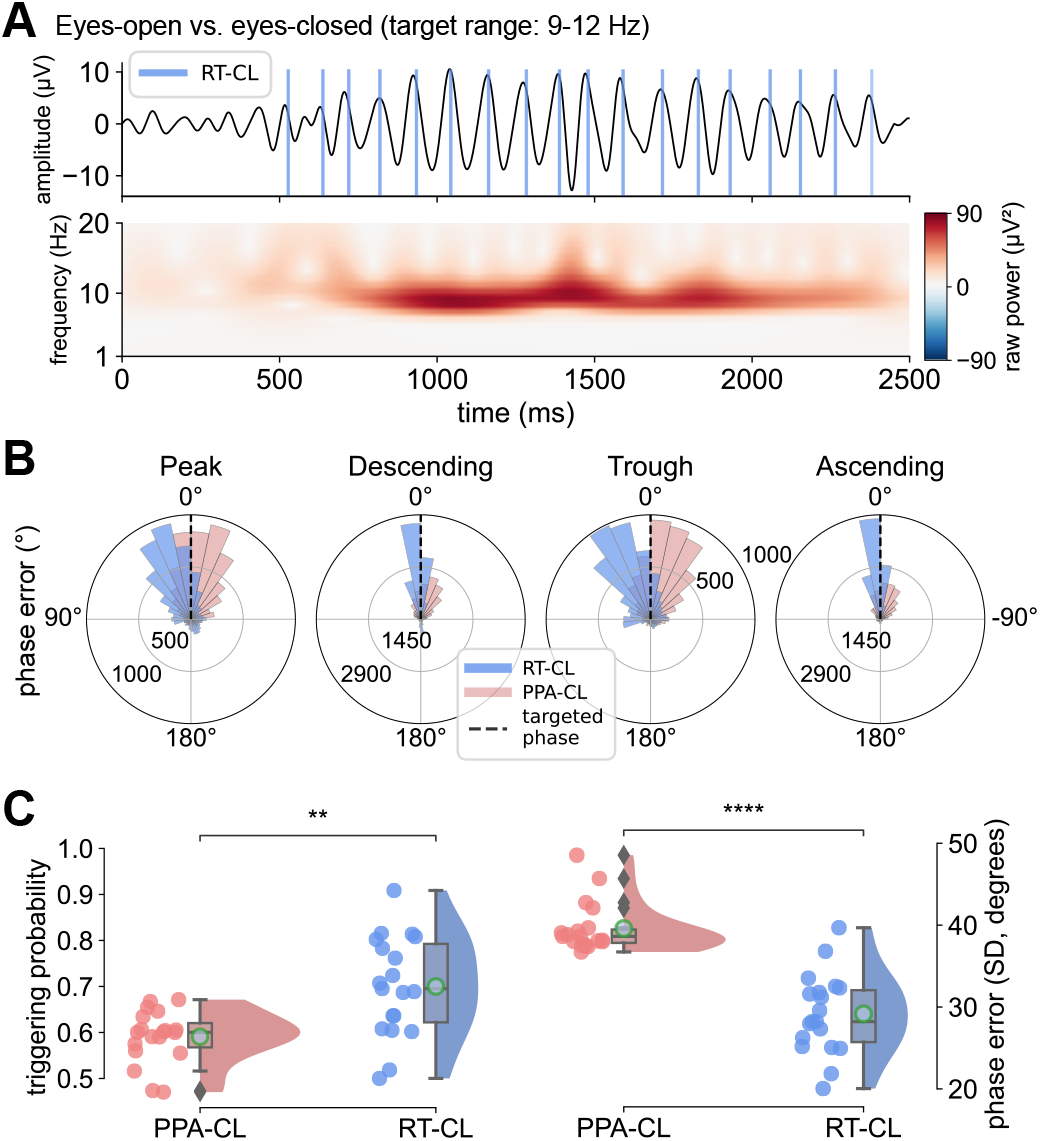
Triggering performance for spontaneous alpha oscillations. **A**: The top trace shows an example of human alpha EEG signal (9-12 Hz) at Oz while targeting peaks using the RT-CL. Blue vertical lines mark the triggering timings. Below is the same time window convolved with a 5-cycle Morlet wavelet showing raw power values, highlighting alpha power increases. **B**: Circular histograms show the distributions of the phase error (deviation of triggered phase from targeted phase) for all subjects and for each closed-loop system (RT-CL: blue, PPA-CL: red) separated by phase target (left to right). Positive values indicate triggering after the target phase occurred, negative values before. Black, vertical lines mark the point of 0° deviation from the targeted phase angle. **C**: Triggering sensitivity (left) and the standard deviation of the phase error (SDPE in degrees, right) for the RT-CL (blue) and PPA-CL (red) systems. Raincloud plots are composed of a density distribution and boxplot for the individual subject values marked by the dots in each scatterplot. Green circles inside the boxplot show the mean value of the distribution and gray diamonds highlight outlier values. Higher triggering probability and lower SDPE indicate better performance. *\*\*: p < 0*.*01, ****: p < 0*.*0001*

#### T-maze RPT oscillations

Fig. 5A depicts a single-trial triggering example and average time-frequency transformation from one subject with RT-CL trigger timings. Consistent with our previous reports [26], [35], robust increases in RPT at right posterior channels (maximal at P8) were detected following feedback presentation. Similar to alpha oscillations, the two systems differed in timing relative to the targeted phase. The RT-CL system delivered triggers shortly after the targeted phase (M_phase error_: 18°), while the PPA-CL delivered triggers before the targeted phase (M_phase error_: -31.5°, Fig. 5B). This temporal difference reflects the RT-CL system’s direct phase detection versus the PPA-CL’s system’s predictive approach. For triggering precision, RT-CL achieved significantly lower phase variability than PPA-CL, *t*(18) = 7.79, *p* = 3.56 × 10^-7^, *d* = 1.84 (Fig. 5B and 5C). The RT-CL system had an average SDPE of 43.2° (SD = 4.3°), compared to 55.1° (SD = 5.2°) for the PPA-CL, representing approximately 12° higher precision. Phase error histograms (Fig. 5B) showed narrower distributions for the RT-CL system, revealing more consistent triggering. Regarding triggering sensitivity, the RT-CL system demonstrated significantly higher triggering probability than the PPA-CL system, *t*(18) = 3.05, *p* = .007, *d* = 0.72 (Fig. 5C, left). RT-CL detected an average of 82.7% of RPT cycles (SD = 13.9%), compared to 72.4% (SD = 9.7%) for PPA-CL, representing a 10% advantage.

**Fig. 5.**
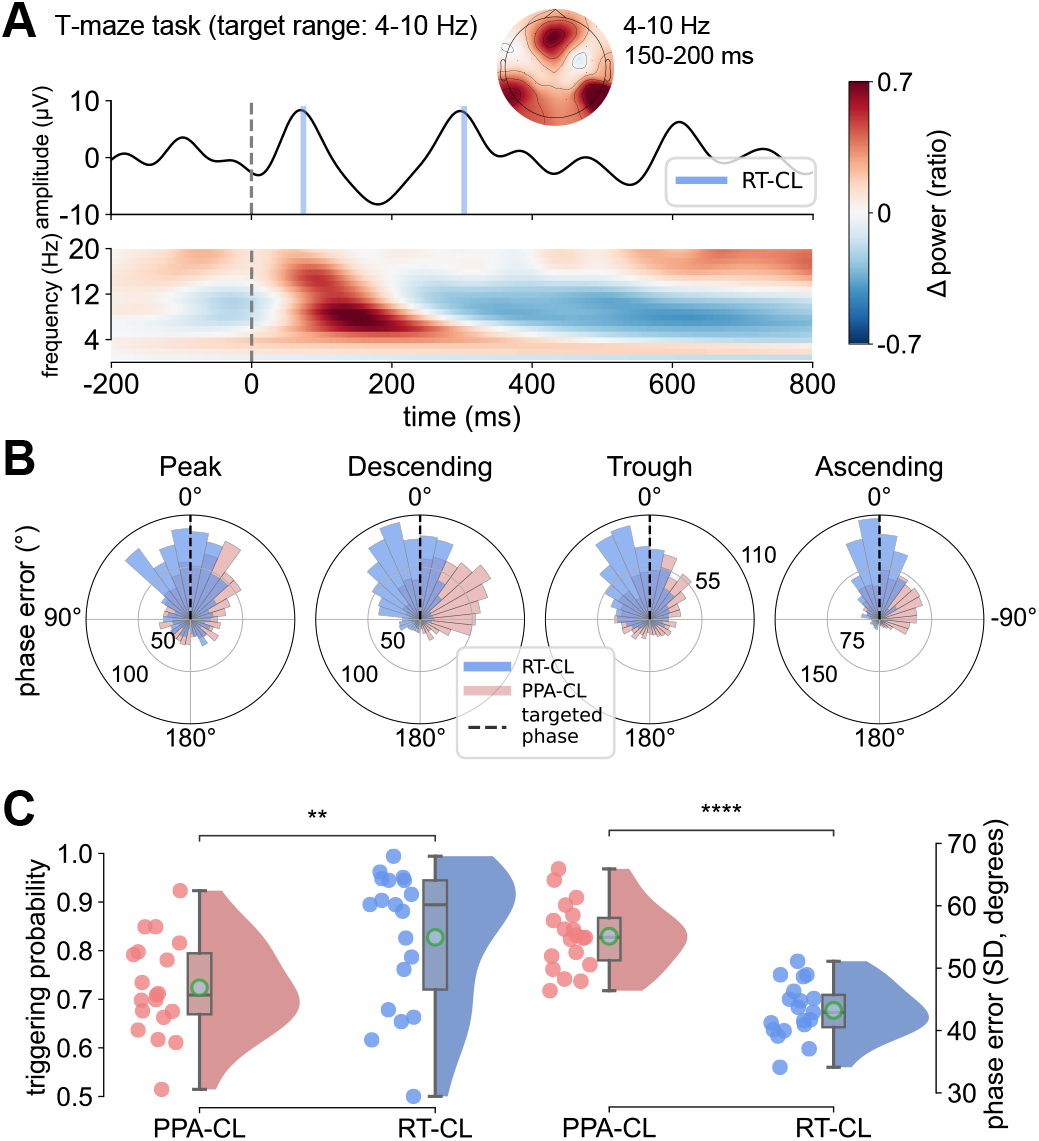
Triggering performance for RPT in the T-maze task. **A**: A single example trial from a subject performing the virtual T-maze task, time-locked to the feedback presentation. Blue bars highlight trigger timings on that trial while targeting 4-10 Hz peaks using RT-CL. Topography (4-10 Hz, 150-200 ms) and spectrogram (channel P8) show the total power results after convolving all trials of the same subject as above with a 7-cycle complex Morlet wavelet and calculating the power change as a ratio of the baseline from -300 ms to -200 ms pre-stimulus. **B**: Circular histograms of the phase error (deviation of triggered phase from targeted phase) for targeting RPT in the T-maze task structured the same as in Fig. 4B (RT-CL: blue, PPA-CL: red). **C**: Analogous raincloud plots to Fig. 4C showing the triggering probability (left) and the standard deviation of the phase error (SDPE, right). As before, higher triggering probability and lower SDPE indicate better performance. *\*\*: p < 0*.*01, ****: p < 0*.*0001*

#### LTM RPT oscillations

Similar to the T-maze task, feedback-related cues in the LTM task elicited robust RPT increases at right posterior channels. A single-trial triggering example with RT-CL and RPT power averaged across trials are shown in Fig. 6A. As in previous analyses, the two systems differed in timing relative to the targeted phase. The RT-CL system delivered triggers shortly after the targeted phase (M_phase error_: 11.5°), while the PPA-CL system delivered triggers before the targeted phase (M_phase error_: -36.9°; Fig. 6B). For phase precision, triggering with RT-CL achieved significantly lower phase variability than with PPA-CL, *t*(18) = 12.59, *p* = 2.32 × 10^-10^, *d* = 2.97 (Fig. 6B and 6C, right). RT-CL showed an average SDPE of 41.9° (SD = 2.8°), compared to 59° (SD = 4.9°) for PPA-CL, representing approximately 17° higher precision. Phase error histograms (Fig. 6B) revealed substantially narrower distributions for RT-CL, with the performance advantage most pronounced in this task featuring continuous movement and greater temporal variability of RPT across events. In terms of sensitivity, the RT-CL system demonstrated significantly higher triggering probability than the PPA-CL system, *t*(18) = 6.89, *p* = 1.93 × 10^-6^, *d* = 1.62 (Fig. 6C, left). The RT-CL system detected an average of 85.4% of RPT cycles (SD = 10.5%), compared to 65.4% (SD = 3.6%) for the PPA-CL system, representing a 20% advantage – the largest sensitivity difference observed across all tasks and frequency bands.

**Fig. 6.**
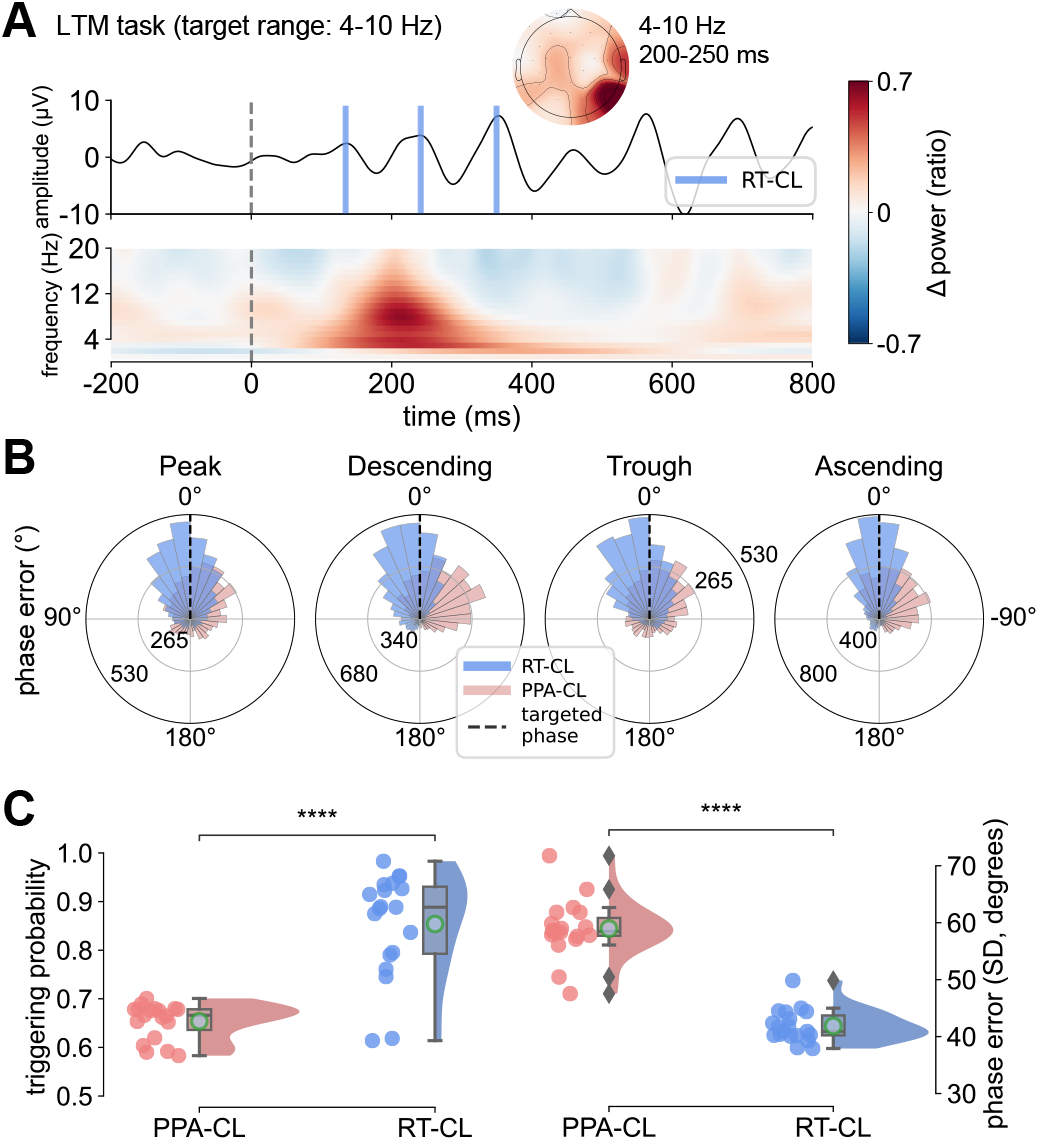
Triggering performance for RPT in the LTM task. **A**: A single example trial from a subject performing the LTM task, time-locked to the feedback presentation. Blue bars highlight trigger timings while targeting 4-10 Hz peaks using RT-CL. The topography (4-10 Hz, 200-250 ms) and spectrogram (channel P8) are the average of all trials after convolving them with the same wavelet as described for Fig. 5A. **B**: Circular histograms of the phase error (deviation of triggered phase from targeted phase) for targeting RPT in the LTM task structured the same as in Fig. 4B and 5B (RT-CL: blue, PPA-CL: red). **C**: Analogous raincloud plots to Fig. 4C and 5C showing the triggering probability and the standard deviation of the phase error (SDPE), with higher sensitivity and lower SPDE indicating better performance. *\*\*\*\*: p < 0*.*0001*

## IV. Discussion

Closed-loop TMS-EEG represents a critical advance for investigating phase-dependent mechanisms during motor, sensory and cognitive processes, and for developing precision neuromodulation interventions. This approach targets the periodic windows of excitability defined by specific phases of ongoing oscillations. Motor-evoked potentials from single-pulse TMS are largest at the trough or rising phase of mu oscillations, with roughly one Hedges’ g separating peak and trough excitability [3]. However, these foundational studies based on phase prediction methods have targeted spontaneous oscillations during rest, and targeting the phase of transient oscillations in real-time during cognitive tasks has yet to be achieved with high precision. The RT-CL system validated here achieves direct phase detection through microsecond-scale hardware processing, enabling reliable targeting of transient event-related oscillations that emerge for only a few cycles during task-relevant processing.

We show that across simulated signals, spontaneous alpha oscillations, and event-related theta activity during spatial navigation, RT-CL demonstrated consistently higher sensitivity (70-85%) and precision (29-43° SDPE) than PPA-CL. When targeting theta cycles in the simulated chirp (i.e., continuously frequency-modulated signal), both systems achieved high performance levels, but only the RT-CL system had consistent triggering performance across the whole theta band. Using PPA-CL on spontaneous alpha oscillations, we detected roughly 59% of alpha cycles with a precision level of 40°, confirming that our results match benchmarks established across multiple laboratories using diverse prediction algorithms for the theta and alpha bands (40-70°, [12], [15], [16], [17], [36]). When using RT-CL, triggering probability was approximately 10% higher and the SDPE was 10° lower, showcasing more consistent triggering on spontaneous oscillations. Finally, we found the most substantial difference between systems when targeting the phases of RPT related to memory encoding. The RT-CL system was 10-20% more sensitive and 12-17° more precise relative to the PPA-CL system, establishing the RT-CL system’s higher precision for testing causal hypotheses about oscillatory phase during active cognitive.

These capabilities of RT-CL versus PPA-CL reflect a fundamental distinction that determines optimal applications for each method. Whereas the RT-CL system depends on fast online detection, phase prediction relies on the assumption that ongoing neuronal oscillations are sufficiently stable to allow accurate phase extrapolation. Other phase prediction systems report lower average phase errors than we achieved, but with larger variability [15], [16], suggesting that on spontaneous oscillations predictions may get closer to the desired phase at the cost of consistency. To date, prediction-based algorithms have demonstrated excellent performance for targeting spontaneous oscillations with high periodicity and signal-to-noise ratio, such as sensorimotor mu rhythms during rest [9], [10], [11], [12], [17]. However, prediction-based approaches face fundamental challenges when targeting EROs that exhibit abrupt phase shifts (e.g., phase resetting) and amplitude fluctuations, precisely the signals implicated in memory encoding, decision-making, and cognitive control.

EROs such as RPT and frontal midline theta, as well as event-related potentials (e.g., N100, N200, P300), have been proposed to arise from partial phase resetting, a stimulus-induced phase shift of ongoing oscillations accompanied by an increase in power [25], [26], [37], [38], [39]. The abrupt phase shift disrupts the temporal continuity phase extrapolation depends on, reducing prediction reliability at precisely the moment when the signal becomes behaviorally relevant. Although novel systems have started addressing this issue by using state space modeling for predictions [17], a more common autoregressive approach avoids theta phase resetting to prevent inaccurate triggering [36]. Beyond phase resetting, the inherent trial-to-trial variability in onset latency and amplitude of event-related potentials and EROs [40], [41], [42] introduces signal variance, complicating prediction accuracy. This variability is reflected in the lower performance of PPA-CL when targeting RPT in the LTM task instead of the T-maze: Due to a single trial in the LTM task involving movement between five locations along a track, RPT may become less stereotyped across events. Thus, only RT-CL maintained strong performance for EROs characterized by trial-to-trial variability. By contrast, PPA-CL performance degraded as predictability decreased.

The mechanistic insights enabled by RT-CL have direct translational implications for precision neuromodulation. Beyond investigating basic neuroscience questions, RT-CL enables a unique approach to clinical intervention: targeting pathological brain states as they emerge during clinically-relevant cognitive or emotional processing. Current FDA-approved repetitive TMS (rTMS) protocols for depression, obsessive-compulsive disorder, and smoking cessation apply rTMS at fixed frequencies independent of ongoing brain activity, yielding response rates of 50-78% [3], [43], [44]. Closed-loop approaches synchronizing stimulation with individual brain states promise to enhance these outcomes by delivering neuromodulation when circuits are maximally responsive. Early clinical trials demonstrate feasibility of phase-specific rTMS synchronized to individualized alpha rhythms, producing EEG alpha band entrainment in depression [45], [46], [47]. Matching the rTMS pulse frequency or inter-burst intervals to EEG oscillations requires continuous oscillations for 2-5 s trains, which is more likely to occur during rest than during cognitive and sensory tasks. Hence, phase prediction approaches may be well-suited for this goal. Conversely, RT-CL specifically enables targeting transient activity during active symptom provocation or cognition – for example, synchronizing TMS with RPT oscillations evoked by stimulus encoding during the LTM task [48]. This “state-and-context-dependent” precision represents a shift from previous closed-loop rTMS trials, with stimulation parameters optimized not only to oscillatory brain state but to the context evoking symptoms.

With novel methods come several considerations that should inform future applications of this approach. RPT during spatial navigation represents a challenging target characterized by a low signal-to-noise ratio and substantial trial-to-trial variability compared to occipital alpha oscillations. While this validates the capabilities of RT-CL under demanding conditions, future work should assess whether the observed performance generalizes to other established event-related EEG signals, such as frontal midline theta during cognitive control tasks [49]. These signals exhibit varying degrees of periodicity and phase consistency, potentially affecting optimal triggering approaches. Additionally, the present validation examined triggering performance without TMS pulse delivery to avoid confounding TMS-induced EEG artifacts. Further research is needed to test whether precise event-related and phase-dependent stimulation with RT-CL produces predicted behavioral and neural effects.

Building on this validation, several applications of RT-CL technology merit investigation. First, targeting higher-frequency oscillations (30-60 Hz gamma) would enable testing phase-amplitude coupling mechanisms proposed to coordinate large-scale networks [50], [51]. While existing phase prediction systems have been successfully used to trigger on faster frequency bands such as human beta rhythms [11], [15], [16], [34], gamma oscillations exhibit pronounced transient burst characteristics potentially unsuitable for prediction-based methods. Therefore, the direct detection of the RT-CL approach may prove particularly valuable for gamma oscillations. Second, implementing complex therapeutic protocols, such as EEG-synchronized rTMS protocols, would advance translational applications. Finally, integrating RT-CL with multi-modal brain state assessment through individualized MRI-based targeting or peripheral physiological markers could enable even more precise identification of optimal neuromodulation windows [52]. As closed-loop technologies transition from research tools to clinical interventions, the demonstrated robustness for transient, variable neural signals positions the RT-CL system as a key technology for the next generation of precision psychiatry and neurology treatments.

## V. Conclusion

The RT-CL compared to the PPA-CL system demonstrated superior and consistent performance for targeting transient EROs through direct phase detection enabled by FPGA-based RT-CL processing EEG data within microseconds. By overcoming fundamental limitations of phase prediction for brief, variable neural signals during active cognition, RT-CL establishes both a methodological foundation for investigating causal phase-dependent mechanisms and a potential platform for developing precision neuromodulation interventions that target pathological brain states during symptom-relevant processing. The capability to reliably synchronize brain stimulation with the fleeting neural dynamics of active human cognition and emotion could open new horizons for understanding brain function and for treating neuropsychiatric disease.

