## Supplementary Table 1 for "Precision phase targeting of event-related oscillations using real-time closed-loop TMS-EEG"

**Table 1. Performance measures by system, task, and phase target**

| Band and Task | System | Target | Sensitivity (%) | Phase Error (M°) | Phase Error (SD°) |
|---|---|---|---|---|---|
| **Alpha (9-12 Hz) eyes-open vs. eyes-closed** | *PPA-CL* | Ascending | 56.4% | -14.3° | 39.4° |
| | | Descending | 59.1% | -13.4° | 38.3° |
| | | Peak | 58.5% | -12.9° | 39.6° |
| | | Trough | 58.3% | -16.1° | 41.0° |
| | *RT-CL* | Ascending | 77.8% | 6.8° | 21.8° |
| | | Descending | 75.9% | 0.0° | 21.1° |
| | | Peak | 69.0% | 20.5° | 35.7° |
| | | Trough | 70.8% | 21.8° | 38.0° |
| **Theta (4-10 Hz) T-maze** | *PPA-CL* | Ascending | 74.3% | -35.7° | 54.6° |
| | | Descending | 72.7% | -46.0° | 56.2° |
| | | Peak | 75.4% | -23.4° | 55.8° |
| | | Trough | 74.0% | -4.3° | 53.9° |
| | *RT-CL* | Ascending | 85.4% | 5.8° | 37.0° |
| | | Descending | 87.2% | 19.8° | 44.7° |
| | | Peak | 89.4% | 7.1° | 47.4° |
| | | Trough | 89.0% | 17.1° | 43.9° |
| **Theta (4-10 Hz) LTM** | *PPA-CL* | Ascending | 66.4% | -36.6° | 60.1° |
| | | Descending | 63.3% | -40.6° | 56.9° |
| | | Peak | 72.1% | -30.3° | 60.6° |
| | | Trough | 65.0% | -21.4° | 58.3° |
| | *RT-CL* | Ascending | 89.1% | 8.2° | 36.2° |
| | | Descending | 87.4% | 22.1° | 40.8° |
| | | Peak | 87.9% | 5.8° | 45.9° |
| | | Trough | 89.1% | 7.9° | 44.6° |
